# OptiDiff: structural variation detection from single optical mapping reads

**DOI:** 10.1101/2022.01.08.475501

**Authors:** Mehmet Akdel, Dick de Ridder

**Affiliations:** Bioinformatics Group, Wageningen University, Droevendaalsesteeg 1, 6708PB Wageningen, The Netherlands

## Abstract

Detecting structural variation (SV) in eukaryotic genomes is of broad interest due to its often dramatic phenotypic effects, but remains a major, costly challenge based on DNA sequencing data. A cost-effective alternative in detecting large-scale SV has become available with advances in optical mapping technology. However, the algorithmic approaches to identifying SVs from optical mapping data are limited. Here, we propose a novel, open-source SV detection tool, OptiDiff, which employs a single molecule based approach to detect and classify homozygous and heterozygous SVs at coverages as low as 20x, showing better performance than the state of the art.

## 1 Introduction

Evolutionary changes in eukaryotic genomes often involve large rearrangements, insertions and deletions that frequently exceed hundreds of kilobases (kb) in length (Y. Li, Zhou, Schwartz, & Ma, 2016; Saxena, Edwards, & Varshney, 2014). These structural variations (SVs) can underlie substantial differences in phenotypes. Large SVs are prevalent between semi-isolated populations of the same species and also appear spontaneously between individuals of a less diverse population. Plant genomes in particular are more prone to the introduction of large SVs, induced by activity of transposable elements present at high frequencies. The evolutionary role of SVs, combined with the agricultural interest in improvement of crop and ornamental plants, has made studying SVs and their phenotypic effects increasingly popular (Gabur, Chawla, Snowdon, & Parkin, 2019; Marroni, Pinosio, & Morgante, 2014).

Over the last decade, two main approaches have been used to detect SVs: microarray-based (Locke et al., 2004; Snijders et al., 2001) and sequencing-based (Kosugi et al., 2019). Microarray-based methods typically have a low resolution and low sensitivity for small SVs. Sequencing-based methods, on the other hand, have been successfully used to detect longer range SVs with high resolution, but are prone to errors at repeat sites and thus are limited in the length of SVs that can be detected (Alkan, Coe, & Eichler, 2011).

Optical Mapping (OM) is an alternative technology to sequencing, which works by labeling high molecular weight DNA at specific sequence motifs and then assembling these single molecule label patterns into a genome-wide map. Optical mapping platforms initially produce image data with 2 channels, where the first channel shows the unspecific backbone labels on the DNA molecules and the second channel shows the specific locations marked by the nicking enzymes or direct label enzyme systems. The first channel is used to determine the border of each molecule, enabling extraction of each molecule while recording the specific labels based on the corresponding pixel coordinates in the second channel. This results in a BNX file which stores molecule information. A single entry in a BNX file contains the length of a DNA molecule, the coordinates of the specific labels within the molecule and the intensity levels of these specific labels. These individual molecules can then be assembled into genome maps, stored in the CMAP format. Entries in CMAP files are based on consensus labels, where each line gives the label ID, coordinate location and the contig ID.

Optical maps can be produced for different individuals and compared to detect large SVs with a higher sensitivity than sequence-based detection methods (Lam et al., 2012). This can mainly be attributed to the technology’s capability of mapping DNA strands of (in principle) any length. However, computational methods for SV detection based on optical map data are scarce. The only commercially available high throughput optical mapping technology is provided by Bionano Genomics (BNG), with the Saphyr (Chan et al., 2018a) as its latest platform. The Saphyr comes bundled with BNG’s closed source software suite and includes two SV detection tools. The first, BNG Solve, takes an assembly-based approach, which involves assembling optical map molecules into consensus genome maps and then comparing these genome maps to a reference genome map to detect SVs (Chan et al., 2018a). This approach is sensitive in an ideal setting where both genome coverage and the N50 molecule length are high. However, for less perfect measurement data the assembly process can yield an incomplete and fragmented genome map, hampering comparison to the reference. This is in particular detrimental to SV detection in plants, as genome sizes are often large and coverage is kept low to keep cost manageable. In addition, DNA extraction procedures can be hard to optimize in plants and often yield sub-optimal material in terms of length due to the presence of DNA-damaging secondary metabolites, such as polyphenolic compounds and tannins (Chan et al., 2018b). BNG’s second tool (Genomics, n.d.) uses a split-mapping approach to detect SVs, and is designed to be used for calling rare variants, thus limiting its application to SVs present in only a small fraction of cells such as cancer SV markers. An open-source alternative to BNG is OMSV (L. Li et al., 2017), which uses a combination of different approaches to call SVs. However, in our hands, OMSV gave good results on the test data provided with it, but yielded very high levels of false positives on our own data.

Here we propose an alternative, open-source SV detection tool, OptiDiff (available at https://gitlab.com/akdel/optidiff), which uses a single molecule segment-matching approach to the reference map to detect and classify SV sites at coverages as low as 20x. To achieve high performance at a wide range of coverages, OptiDiff uses a reference molecule set to obtain background mapping levels in all genomic regions on the reference. It then calculates the ratio between this background mapping rate and the SV candidate molecules’ mapping rate to detect SV sites. Based on this segment-match information, OptiDiff then applies a simple rule tree to classify the type of structural variation. We compared OptiDiff to the latest software from BNG (BNG Solve) for SV site detection and classification on simulated data. We also used real optical mapping data from the Cvi1 accession of *Arabidopsis thaliana* and called SVs with respect to the reference accession (Col0). Performance was measured in terms of overlaps between SV calls and the genome alignment of the near-chromosome level assembly of Cvi1 (Jiao & Schneeberger, 2019) to Col0 (TAIR 10). OptiDiff overall was superior in terms of precision and recall in the simulation settings and showed a higher level of consistency with the Cvi1-to-Col0 genome sequence alignments. Moreover, since OptiDiff works with single molecules instead of generating an assembly, it is an order of magnitude faster than the software offered by BNG. However, OptiDiff requires more optimisation in terms of the preliminary conversion of BNX data into OptiDiff’s compressed molecule representation, as the variability on BNX molecule quality can potentially have an effect on this process.

## 2 Methods

### 2.1 Overview

The input to OptiDiff is an optical map molecule set (BNX) of a potential variant genome, an optical mapping based reference genome map (CMAP) and a molecule set (BNX) of the corresponding reference genome. It then uses mapping patterns derived from matching molecule segments (referred to as segment-matching) to the reference genome map. Some areas of the genome may have a higher or lower coverage than average, due to local chromatin characteristics. In order to correct for this coverage variation, a background profile is first created by segment-matching reference molecules to the reference genome map. Then, the SV candidate under investigation is segment-matched in the same way and the pattern of differences in their profiles is used to detect structural variation (Figure 1).

**Figure 1:**
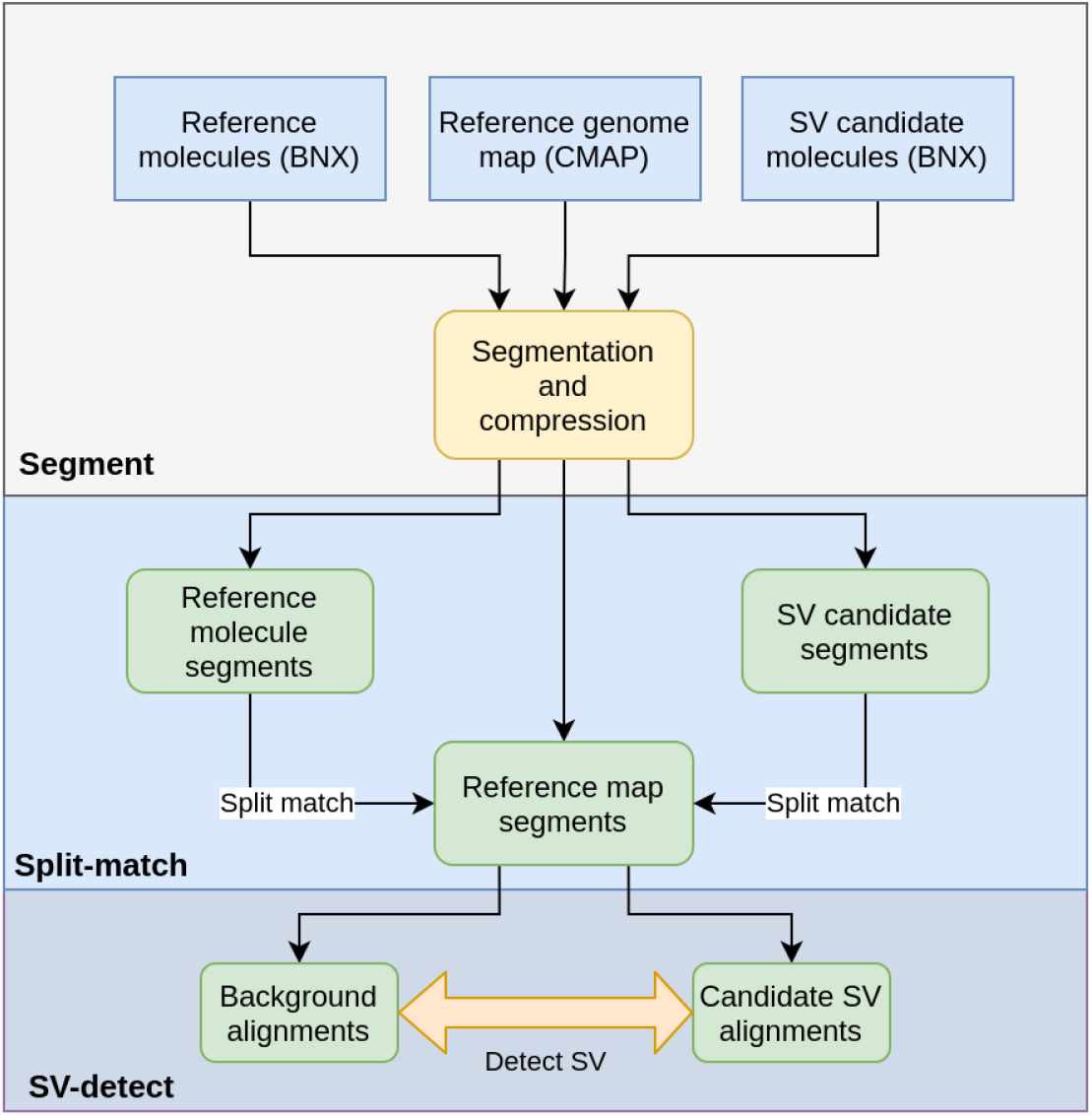
Overview of the OptiDiff SV detection algorithm, divided into three phases: molecule segmentation, segment-matching and SV site detection. An overview of SV type classification is shown in Figure 4.

### 2.2 Molecule segmentation

The first phase of the algorithm involves segmentation of molecules and the reference map. Molecule input data (both candidate SV and reference molecules) should be supplied in the BNX file format and the reference genome map in the CMAP format, as provided by BNG Solve. Molecules are segmented by splitting into regions of a preset length *s* (default *s* = 138 kb), starting at each label site (in the forward orientation relative to the channel). Segments which start with labels closer to the end of the molecule than the set segment length are discarded. As a result, molecules are described by a list of segments in the order of the labels used as split points. For the *in-silico* reference map, each chromosome is similarly converted to a list of segments, this time in both the forward and reverse orientations. Segments are subsequently represented as digital signals, in which each unit corresponds to a DNA region of length *l* and each peak is represented by a square wave (with arbitrary amplitude) *p* units wide. Unit length l and peak width *p* were set based on our previous analysis of raw optical map signals (Chapter **??**) and on BNG imaging specifications (*Optical Mapping - Saphyr Whole Genome Imaging*, 2021) as l = 500bp and p = 10. The digital signal thus represents measurements at the maximum label resolution that the BNG equipment can achieve. However, the signals are often sparse and contain little information for long stretches of DNA. This makes matching long molecules to the reference genome map based on these signals not computationally efficient. Therefore, as a last step, we compress each segment into a binary string of a given number of bits (64 bits by default) to be used in the matching step, through linear interpolation. In this compressed segment, a 1 represents a label site in the original signal and a 0 its absence (Figure 2).

**Figure 2:**
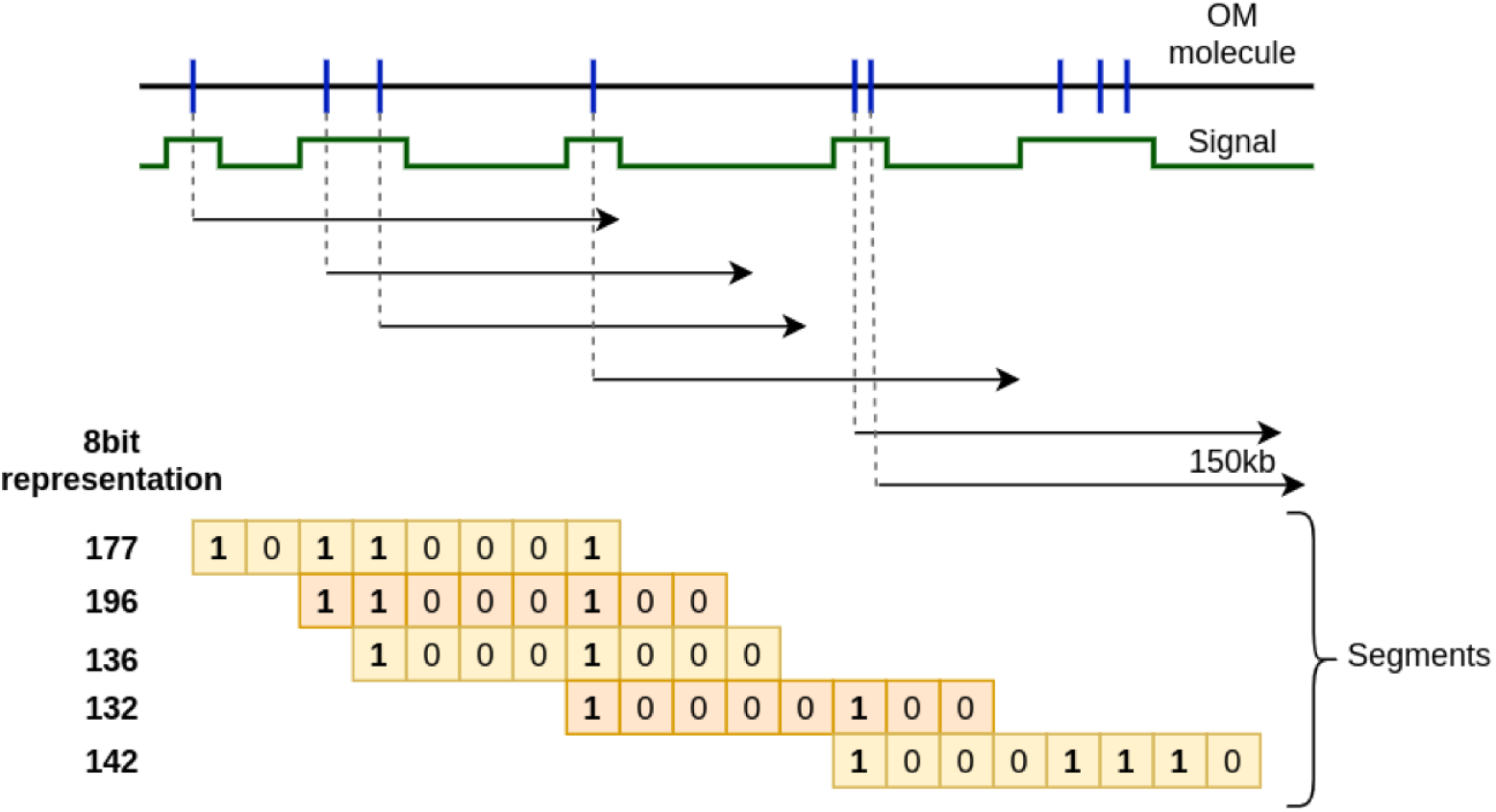
A molecule, represented by a square wave signal, is split into 5 segments. In this example each segment is compressed to a string of 8 bits (the actual algorithm uses 64 bits). Each label site is stored as a 1, each background site as a 0.

### 2.3 SV detection and classification

The SV detection process starts with matching molecule segments, from both molecule sets, to reference segments.

Reference segments are ordered according to the genome location of their start label. Molecule segments are similarly ordered and then matched to the reference according to their bit-wise Hamming distance to each reference segment, with a distance threshold *t* dictating a positive match. This threshold depends on the number of labels *n_r_* which fall into a reference segment *r*, i.e. 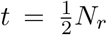. This corresponds to the fact that the number of errors (false positive and negative peaks) correlates with the number of peaks rather than the segment length. The segment length s was chosen to be 138 (as explained below) to have a specific enough segment to appear on average once throughout in the genome and thereby avoid multiple matches as much as possible. In case of multiple matches, we randomly choose one position.

#### 2.3.1 Segment-match patterns

In principle, SVs can be detected immediately as patterns in the segment-matching results. A molecule with any type of SV results in an alteration of segments which start at label sites covering the SV site. These segments become more difficult to match to the reference, leading to gaps in the segment-matching pattern (see Figure 3). Single molecule segment-matching patterns that form the basis for SV detection algorithm are:

1. **Large deletion**: deletions, with one or more missing segments, result in a gapped segment-matching pattern with both altered (due to shifts in the pattern) and missing segments.
2. **Inversion**: since reference segments are created for both genome orientations, molecule segments are matched to both forward and reverse reference segments and the orientation of the match is recorded. An inversion larger than the segment length results in at least one segment matching in the opposite orientation to the others.
3. **Translocation**: molecules which overlap a translocation boundary contain at least one segment of the original sequence and at least one segment of the translocated sequence, if both sequence sections are larger than the segment length.
4. **Small SVs or insertions**: a segment-matching pattern with gaps which cannot be assigned to one of the types above due to its size being smaller than the segment length, or an insertion.

**Figure 3:**
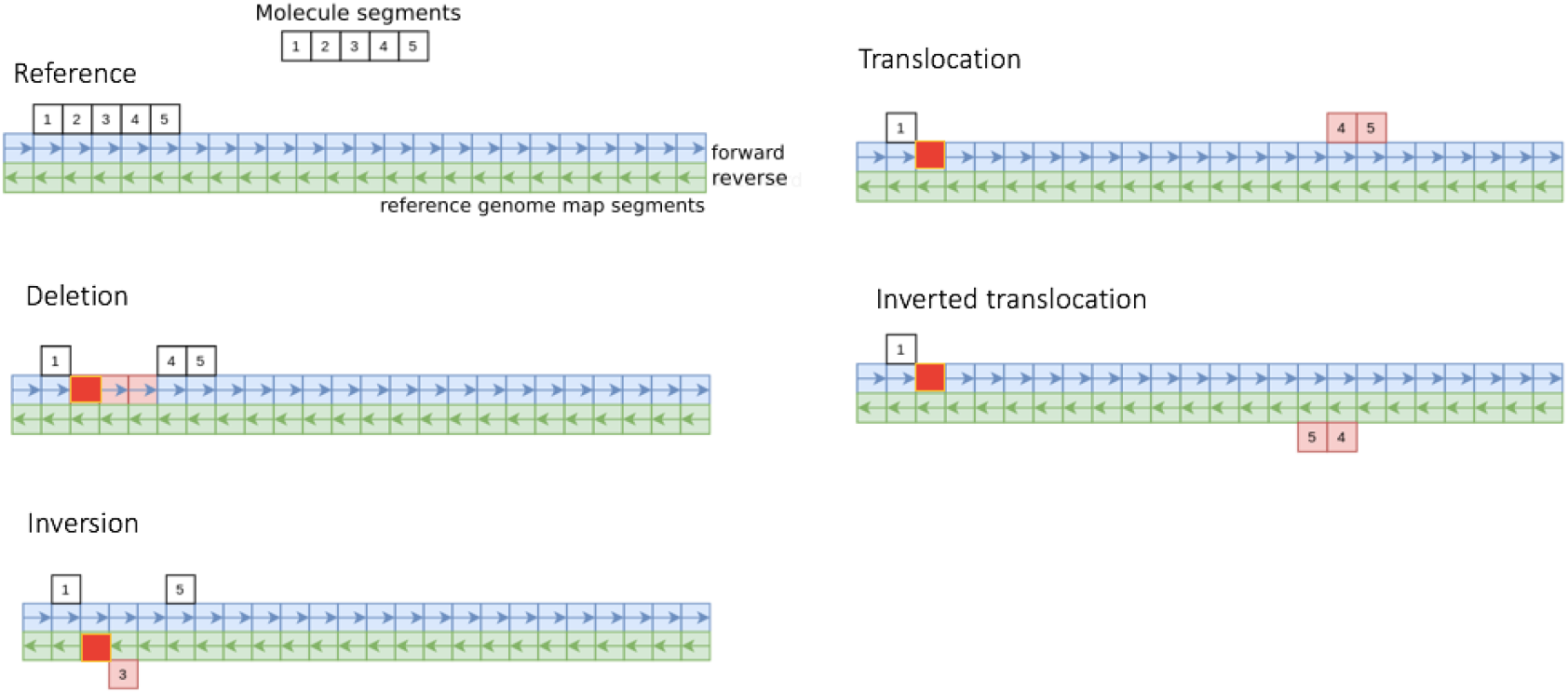
Segment-match patterns that correspond to different types of SV. The “forward” and “reverse” segments represent forward and reverse orientations of the reference genome map. The bright red boxes represent the mismatch segment found in common between all types of structural variants.

However, we cannot simply use single molecules’ matching patterns as direct evidence for SVs, as optical mapping data is often noisy (high mislabeling frequency, DNA stretching and image quality issues) and results in false positive/negative matches. Thus, we instead look at the consensus of all molecules mapping to a region with similar SV patterns as a basis for detection and classification of SVs. We do this by initially generating segment-match coverage profiles for both background and SV candidate molecule sets, with a 500bp resolution. After segment-matching all molecules in a given set onto the genome map, we assign and store matching molecule IDs to each reference genome segment in forward and reverse directions. This information is used for two purposes, to 1) detect SV locations and 2) classify the type of SV at a detected location. For detection, we then create segment-match coverage signals for both molecule sets. For each molecule segment, we add a coverage of 1 starting from the label coordinate of the matched segment on the reference, *m,* up to *s* + *m.* Coverage signals for both background and SV candidate molecule sets are generated by repeating this process for all matching segments. Finally, we normalize these coverage signals by using a robust scaling algorithm (implemented in the Scikit-learn Python library (Pedregosa et al., 2011)), using only values between the 15%th and 85%th percentiles to estimate scaling parameters, to prevent the effects of outliers in low coverage molecule samples (Figure 5(a)).

#### 2.3.2 Detection of SV locations

For detection of SV sites, we obtain the squared log difference (*SQLD*) between the normalized reference segment-match coverage signal *R* and the candidate coverage signal *C*:

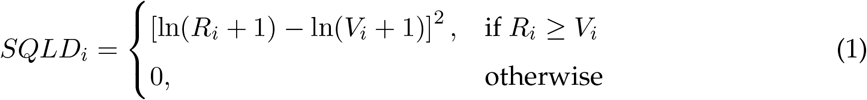

We log-transform the coverage signals to reduce the effect of fluctuations in difference in high-coverage regions (usually caused by repeat elements) and increase the contribution of differences in low-coverage regions. We can then detect peaks in this coverage signal which exceed a given minimum signal-to-noise ratio (SNR), calculated as the peak height divided by the median height of the coverage signal. We used a minimum SNR of 35 throughout this study to filter out low peaks, as these are more likely to be false positives. The resulting detected peaks correspond to low coverage regions in the candidate signal compared to the background, as illustrated in Figure 5(b). The genomic location of the peak and the molecules in the peak region forms an *SV seed,* i.e. the starting point to subsequently classify SVs into specific types (Figure 4).

**Figure 4:**
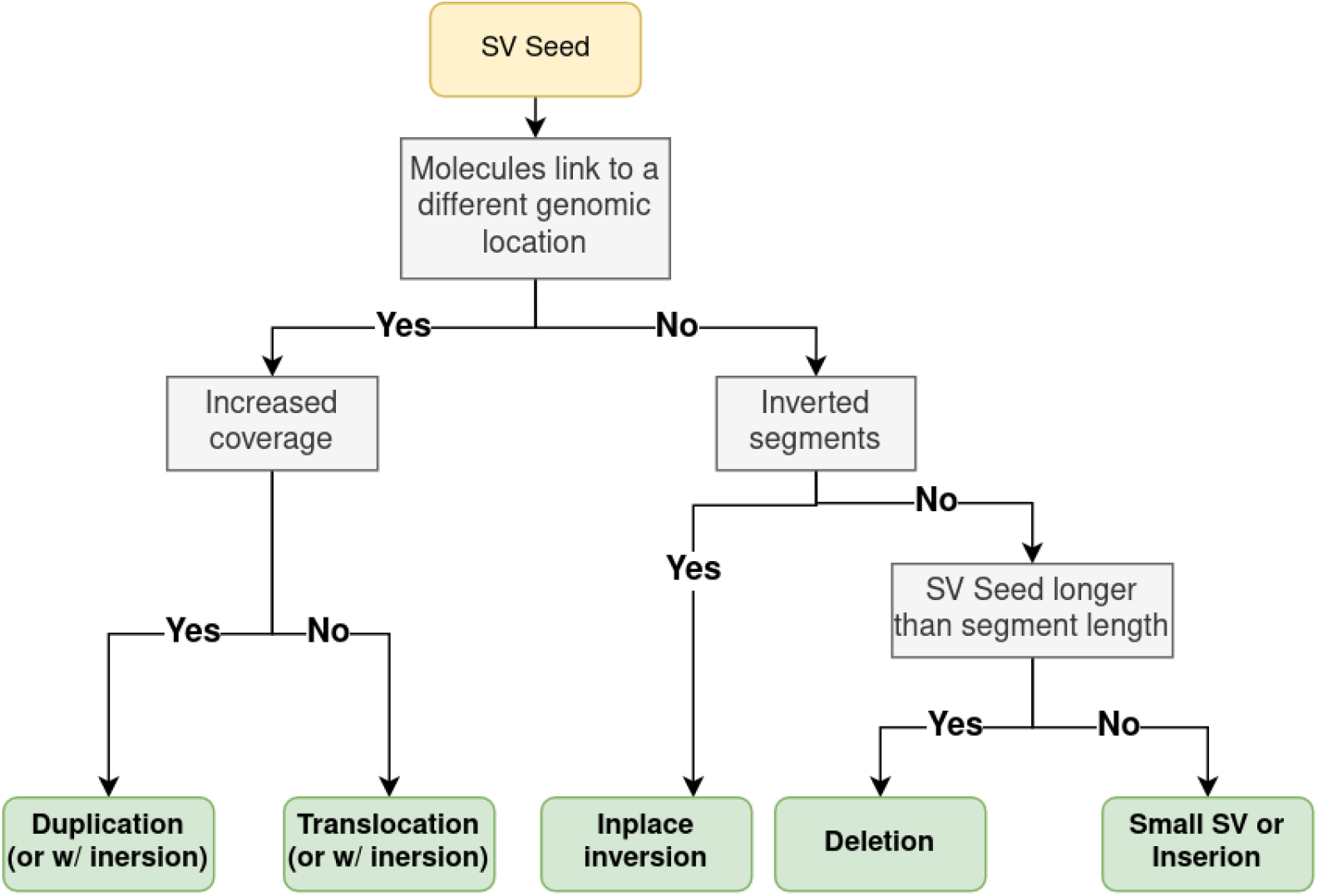
Flowchart of decisions taken to classify different SV types.

**Figure 5:**
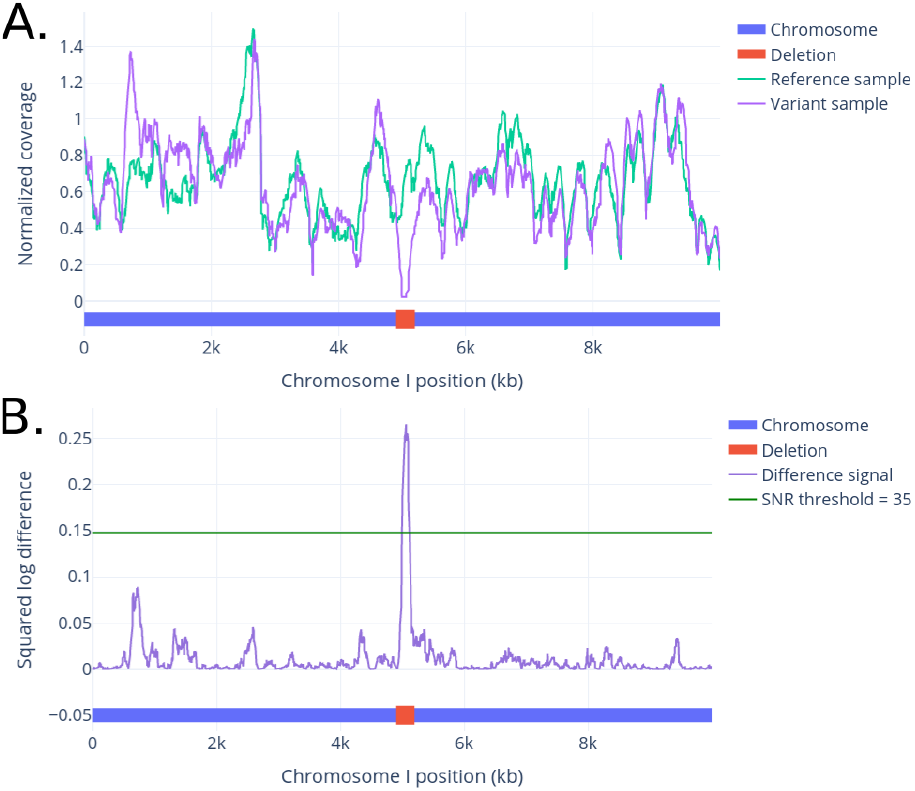
(A.) Coverage signals of a simulated reference genome and a variant containing a deletion, within 10Mb distance from the start of Heinz tomato chromosome 1 (Hosmani et al., 2019). (B.) An SQLD peak at the SV site is easily detected by considering the ratio between the two coverage signals.

#### 2.3.3 Translocation and duplication classification

Classifying an SV seed as a translocation or a duplication relies on creating signals, similar to the segment-match coverage signal, using only matched segments of molecules overlapping in and around the SV site. This process is performed on both the SV candidate and reference molecules. The algorithm is as follows:

A. Reference molecule segments overlapping the beginning of the SV location *x* up to segment length *s* upstream are retrieved (Figure 6A, left).
B. Reference molecules are found which have at least one segment matching with any of the retrieved reference molecule segments, to create a reference coverage signal based on all segments in these molecules (Figure 6B and 6C, left).
C. This process is repeated for the candidate molecules to obtain a candidate coverage signal (Figure 6A-C, middle and right).
D. The candidate signal is divided by the reference signal and the highest peak in the resulting signal above an SNR of 4 is found. This peak marks the start site of the translocated/duplicated sequence (Figure 6D).
E. Steps A-D are repeated downstream of the end of the SV location, *y,* up to the segment length s, and the resulting peak marks the end site of the detected translocated/duplicated sequence (Figure 6E).
F. Using the start and end sites detected above, it is determined if this region is a translocation or duplication. Duplications are marked by a significant increase in mapping coverage of the region between the detected boundaries compared to the rest of the reference (Figure 6(c)). First, the ratio between the candidate and reference coverage signals is computed. A *t*-test between ratios (in 500bp blocks) inside the detected area and outside that area (i.e. the rest of the chromosome) is then performed. If the candidate region has a significantly higher mean ratio (p < 0.001) then it is classified as a duplication, otherwise as a translocation (Figure 6F).

**Figure 6:**
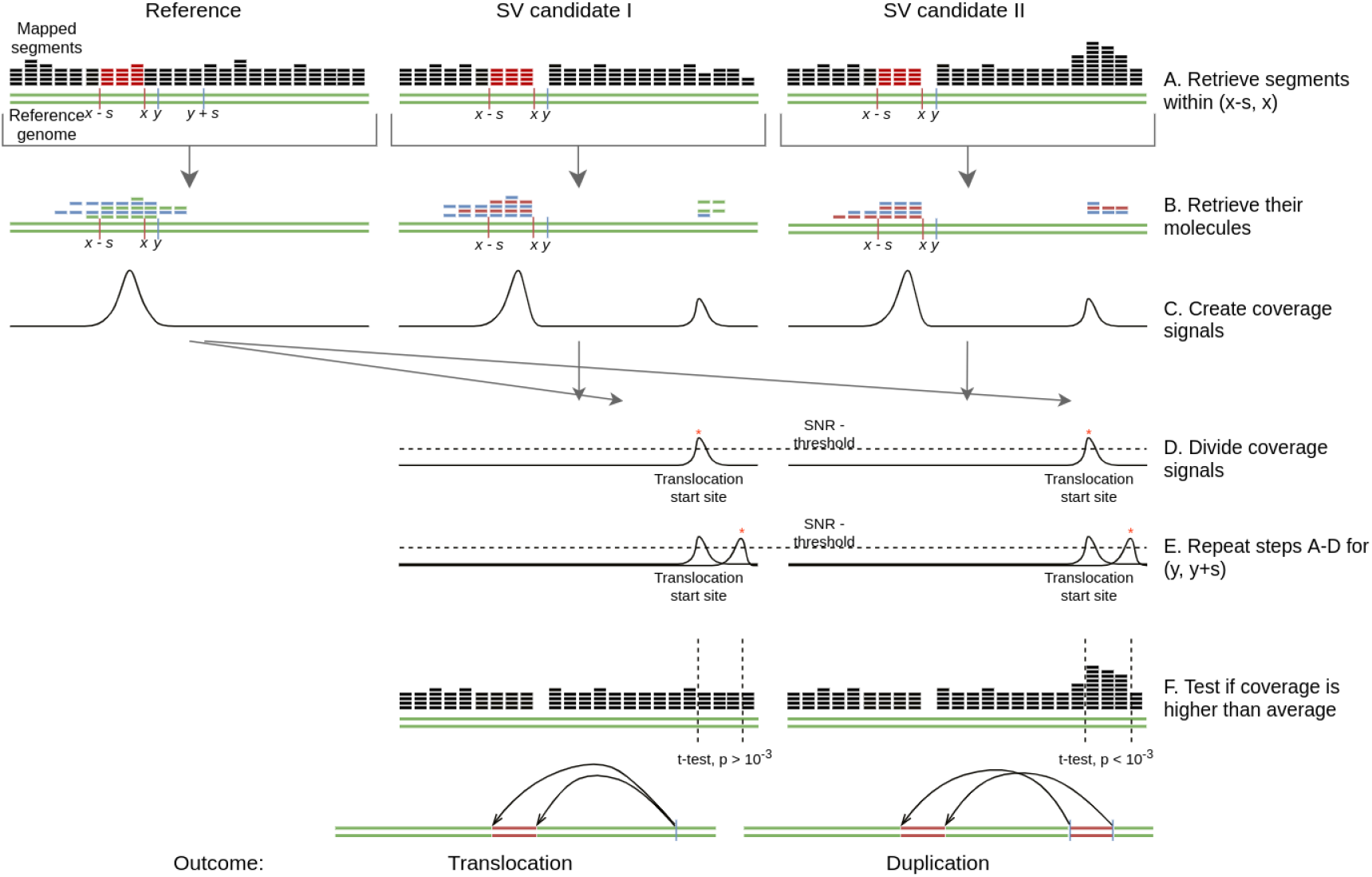
The translocation and duplication classification workflow. After detecting an SV location between coordinates *x* and *y,* these steps are followed: (A.) All segments within *x* and *x* – *s* (where *s* is the segment length) are retrieved for both reference and SV candidate data. (B.) The remaining segments from the molecules which overlap with segments from the previous step are obtained and used to create a coverage signal. (C.) The resulting signal is divided by the reference counterpart. (D.) A peak corresponds to the translocation start coordinate. (E.) The process from A.-D. is repeated on a downstream *y* + *s* section of the genome map to obtain the translocation end coordinate. (E) The coverage of the translocated region is tested to decide on duplication or translocation.

A limitation of this algorithm is that only translocations/duplications longer than the segment length can be detected.

#### 2.3.4 Inversion classification

Similar to translocations and duplications, molecules at inversion boundaries also show a specific pattern, i.e. two subsequent segments matching in opposite orientations (Figure 6(d)). We make use of this shift in segment orientations and compare the frequency of occurrence to that in reference molecules to pick out inversions:

A. As before, reference molecule segments overlapping the beginning of the SV location *x* up to segment length *s* upstream are retrieved (Figure 7A, left).
B. Reference molecules are found which have at least one segment matching with any of the retrieved reference molecule segments, and which match in the opposite direction to the retrieved segments (Figure 7B, left). These are used to create a reference signal (Figure 7C, left).
C. This process is repeated for the candidate molecules to obtain a candidate coverage signal (Figure 7A-C, right).
D. The candidate signal is divided by the reference signal and the highest peak in the resulting signal above an SNR of 4 is found. This peak marks the end site of the inversion sequence (Figure 7D).
E. The above steps are repeated downstream of the end of the SV location, *y,* up to the segment length *s*, and the resulting peak marks the start site of the detected inversion sequence (Figure 7E).

**Figure 7:**
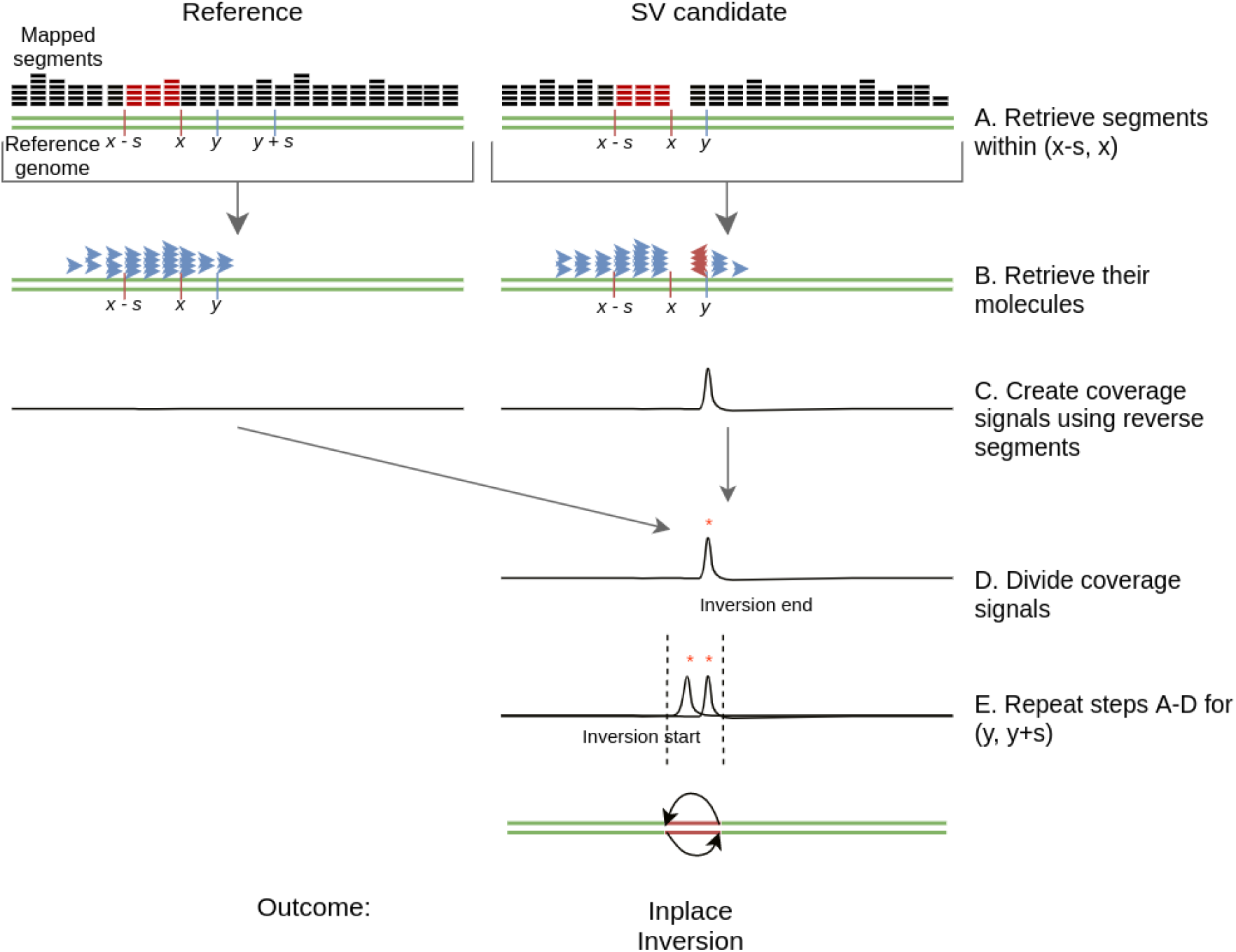
The inversion classification workflow. After detecting an SV location between coordinates *x* and *y*, these steps are followed: (A.) All segments within *x* and *x* – *s* (where *s* is segment length) are retrieved for both reference and SV candidate data. (B.) The remaining segments from the molecules which overlap with segments from the previous step are obtained; only those matching in reverse orientation are used to form a coverage signal. (C.) The resulting signal is divided by the reference counterpart. (D.) A peak in the signal corresponds to the inversion end coordinate. (E.) The process from A to D is repeated on the reverse strand of the genome map to obtain the inversion start coordinate.

It should be noted that if the SV site was not also classified as a duplication or translocation, then the inversion is classified as an inplace inversion. Otherwise, the inversion is classified as a translated inversion or translated duplication. As for translocations and duplications, only inversions longer than the segment length can be detected.

#### 2.3.5 Deletion detection

When translocation, duplication and/or inversion detection return negative results, we are left with deletions, unclassified short SVs and insertions of any length as possibilities. To distinguish between these choices, we look at the width of the SV seed peak (calculated as the width obtained on extending the peak site in both directions until an SNR of 1 is achieved). If the width is larger than the segment length, we categorize the SV seed as a deletion. Otherwise it is either an insertion of any length or another SV which is too short to classify. With the current approach, it is not possible to distinguish between these.

## 3 Results and discussion

### 3.1 OptiDiff detects SV sites with high specificity

Detection of SV locations with a high level of confidence is challenging, yet a prerequisite to successful SV classification. OptiDiff accomplishes this by labeling putative detection sites as SV seeds (as described in Section 2.3.2). Our simple SQLD approach makes this initial identification robust even in low coverage situations. To measure OptiDiff’s performance in this task, we simulated four types of SVs (deletion, inplace inversion, translocation and duplication) of 300kb length at random locations and compared OptiDiff’s results to those obtained using the BNG Solve software. The simulations were based on a 50Mb fragment at the start of chromosome 1 of the tomato genome. We simulated genome maps of 20 random SVs for each SV type. Based on these genome maps, we used OMSim (Miclotte et al., 2017) to simulate optical mapping molecules with realistic noise profiles at coverages ranging from 20x to 120x. In our evaluation procedure, we mark a detected SV as a true positive if it overlaps (by any length) with the simulated SV.

Results are shown in Figure 8, where the red and blue areas together indicate true detections and the purple area indicates false detections. We find that OptiDiff is highly sensitive towards detection of SVs at coverages above 60x, with a low false detection rate in all types of SVs except duplications. Translocations and deletions are similar in terms of the initial detection phase, as they both involve parts missing with respect to the reference. However, duplications can only be detected by spotting the inserted site, as the duplicated part stays intact. Even though OptiDiff’s precision is higher than that of BNG Solve, the false detection rate does not seem to decrease with higher coverages for either of the tools. BNG Solve shows higher numbers of false detections (nearly 2x the number of OptiDiff) for all types of structural variation, inplace inversions being the highest. BNG Solve performed worse at finding SV sites within true duplication events above 40x coverage. A single duplication introduces a long repeat, which can cause complications in the assembly process as only the molecules longer than the repeat can help obtain a complete and correct assembly. Since, unlike OptiDiff, BNG Solve uses fully assembled contigs to detect SVs, increased assembly errors could explain its reduced SV detection performance.

**Figure 8:**
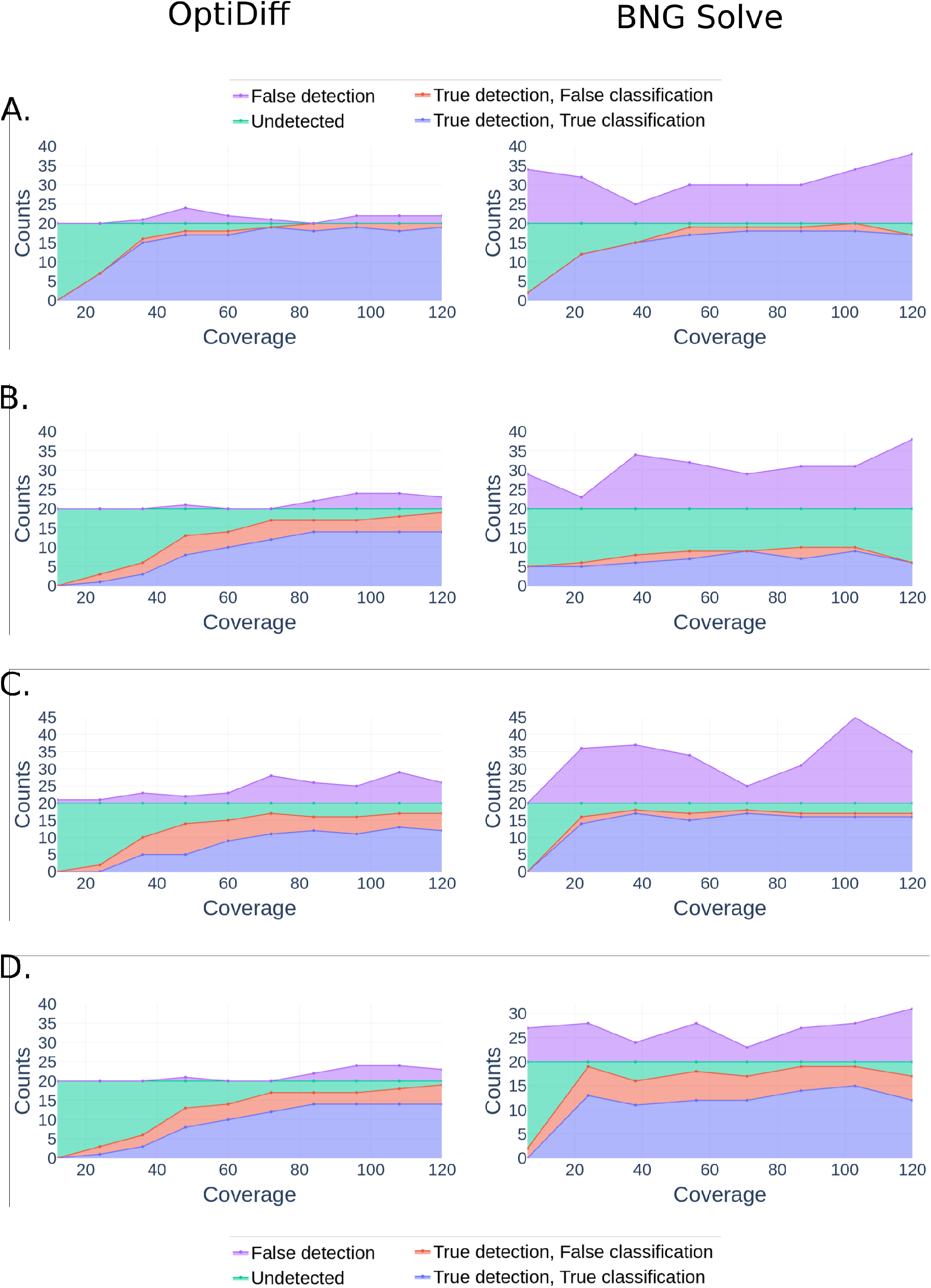
SV detection and classification rates per SV type, for OptiDiff and BNG Solve, for coverages between 20x and 120x. The four types of SVs are A. deletion, B. duplication, C. inversion and D. translocation. Each SC type is simulated 20 times, representing the maximum possible number of true detections and classifications. The purple area above 20 indicates the false detections made outside the simulated SVs. The cyan area shows the false negative detections, red the true detections with false classifications, and blue the correctly classified true detections.

**Figure 9:**
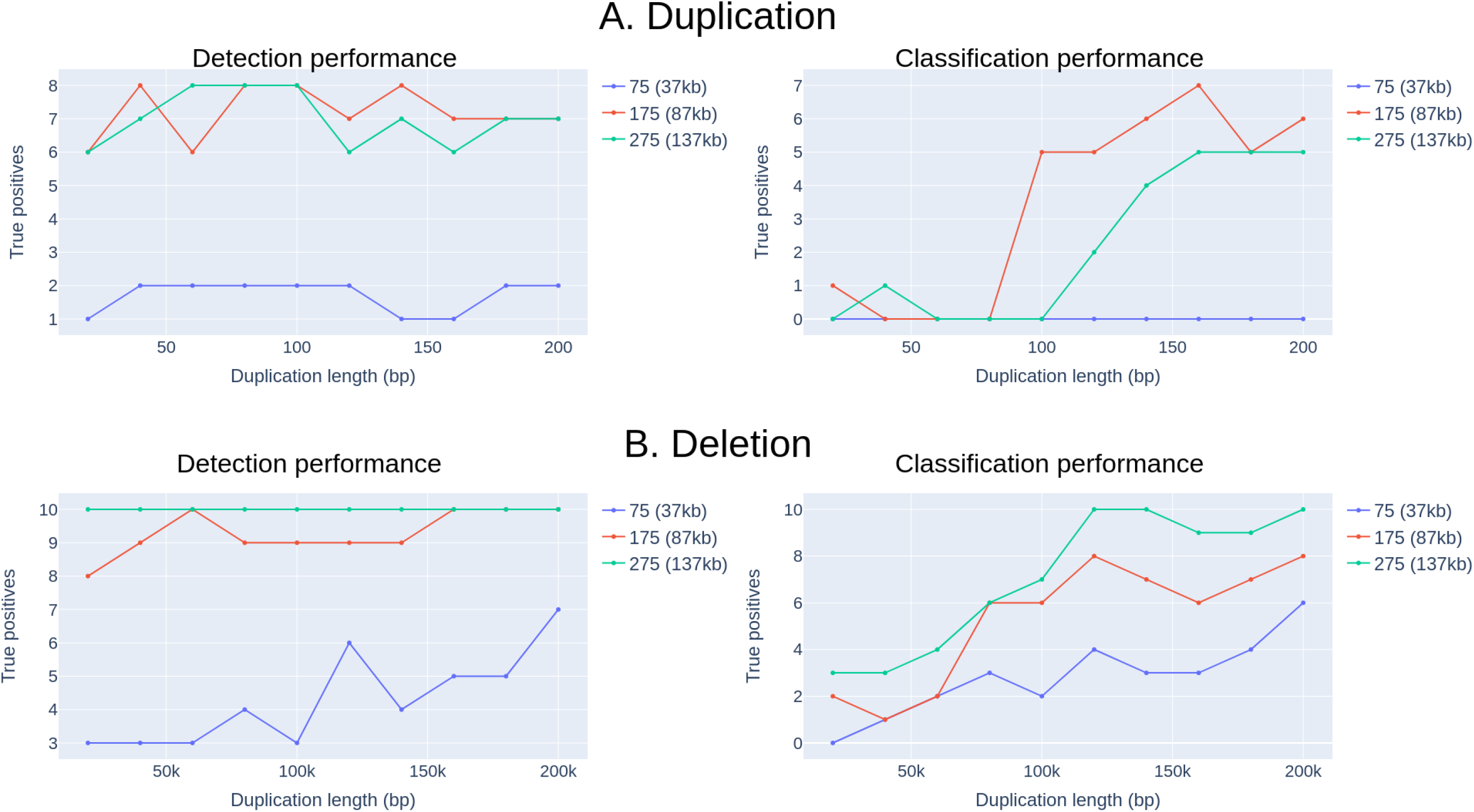
Detection and classification performances for OptiDiff on simulated duplications (A.) and deletions (B.) with lengths ranging from 20k to 200k basepairs. The simulations are repeated using 10 randomly selected sites within the 50Mb chromosome; shown are the number of true positives recovered for each SV length. Three different segment lengths were compared (see legend).

### 3.2 OptiDiff outperforms BNG software in classifying duplications

In contrast to SV detection, SV classification is a more complicated task that requires numerous long molecules with informative segment-match patterns. Here we test OptiDiff’s SV classification capabilities and compare our results to those obtained using BNG Solve, with the same simulation data as used in the previous section. The results are also shown in Figure 8, with blue representing correctly classified SVs and red representing correctly detected but falsely classified SVs. A major difference between the two methods lies in the classification of duplications, where BNG Solve misclassifies all detected duplication events as translocations. Another apparent difference is in inplace inversion classification performance, where BNG Solve performed better overall. At low coverages (< 60x), BNG Solve performed better at translocation classification, although the results of the two tools are similar at higher coverages. Lastly, deletion classification performance was comparable for both tools at all coverages. Overall, given the similar classificaton performance but improved detection performance, OptiDiff is a good alternative to BNG Solve.

### 3.3 Trade-offs in the classification performance of duplications and deletions by altering segment length

Throughout this study, we chose to set the segment size to 275 units (138 kb), which provides a good overall performance. Lowering the segment size allows OptiDiff to detect smaller SVs, but simultaneously makes it progressively harder to classify detected SVs. Here, we test OptiDiff’s limits in detection and classification of short duplications and deletions, by simulating SVs of shorter lengths (9). As the minimum length needed for classification is the segment length, we tried a range of smaller segment lengths (37, 87 and 138 kb). Duplication classification improved for short duplications when lowering the segment size (from 138kb to 87kb) without losing detection performance, although this trend does not carry on to 37kb segments. Since duplication classification requires a label pattern to fall into the duplication site, its performance will suffer from lowering segment length. However, this trade-off is not seen for deletions. This is due to the decreased uniqueness of label patterns in short segments, which more easily match multiple locations on the genome. This can result in a loss of segments through unspecific matching, and increases background noise which becomes detrimental to the SV detection and classification algorithms. Taken together, for increasing duplication classification performance, lower segment lengths are favoured while the opposite is true for deletions. It should be noted that our algorithm labels short SVs as unspecific SVs as these can also be insertions.

### 3.4 OptiDiff can detect and classify heterozygous SVs

The results above demonstrated the use of OptiDiff for detection of homozygous SVs. For the algorithm underlying OptiDiff, heterozygous SVs are harder to detect, largely due to the fact that the effective coverage depth is half of that of homozygous sites. To investigate performance on heterozygous sites, we simulated the four types of SVs as before but at a high coverage setting (240x) that can be achieved with a single flow-cell. We then assessed detection performance in the same way as before. The results are shown in Figure 10. OptiDiff classification results are comparable in deletion events, outperform BNG Solve in duplication classification, and underperform in calling inversions and translocations at coverages below 80x. Overall, the detection and classification performance for both tools has decreased compared to homozygous SVs.

**Figure 10:**
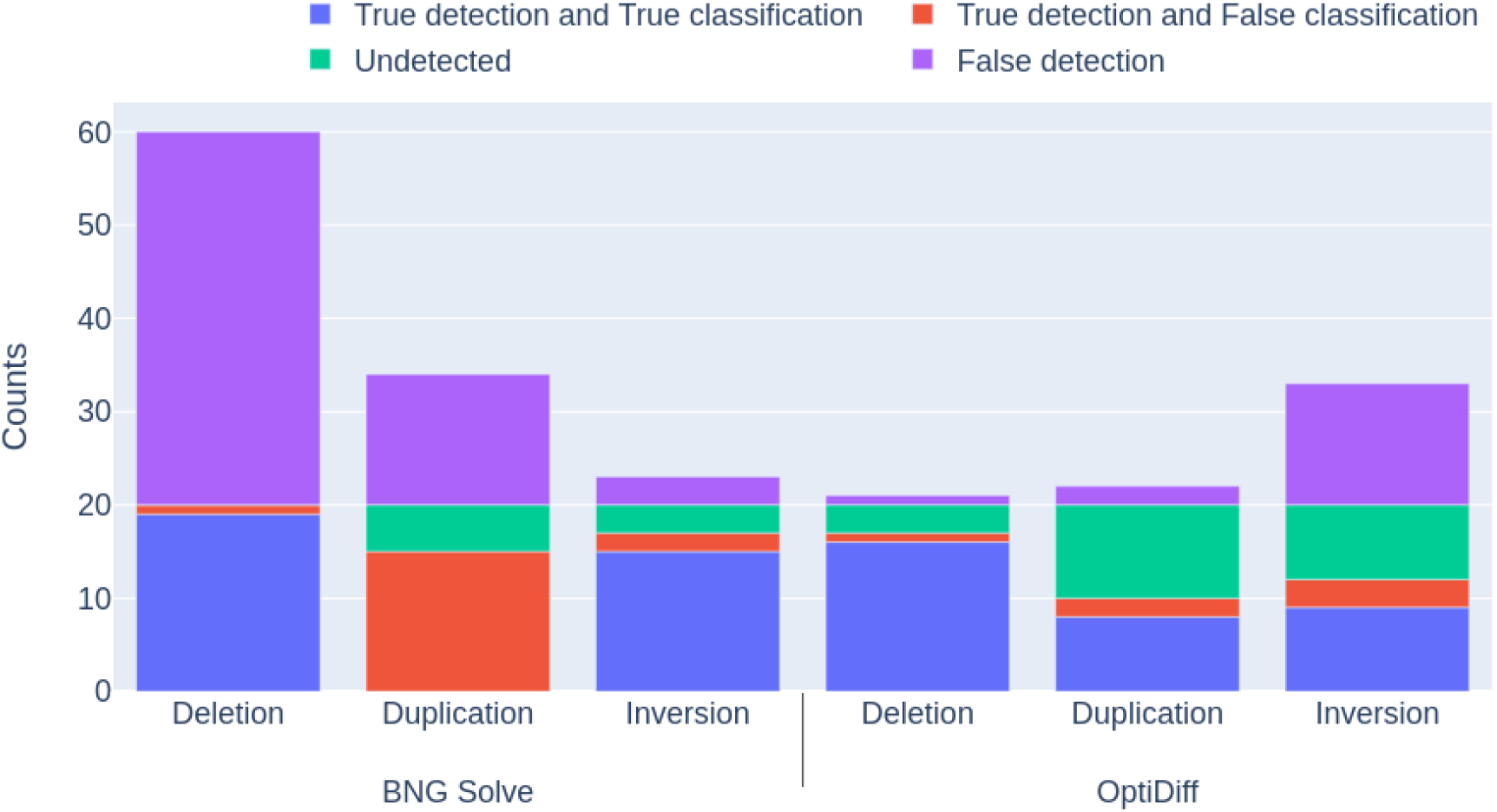
Heterozygous SV detection and classification performance using BNG Solve (left) and OptiDiff (right), at a coverage of 240x. 20 of each type of SV are simulated at random genomic locations.

### 3.5 Detected deletions correspond to gaps in whole genome sequence alignments

In an application on real-world data, we detected structural variations in optical mapping data obtained from two different accessions of *Arabidopsis thaliana* (Kawakatsu et al., 2016). In contrast to the simulated datasets above, where the structural variant sites represented the only points of difference, here there is additional sequence variation which can result in lower mapping performance. We performed whole genome sequence alignments of the near-chromosomal level Cvi genome sequence with the reference genome sequence Col-0 (TAIR 10) (Jiao & Schneeberger, 2019) using MUMMER and extracted gaps longer than 15kb in the alignment. Similarly, we used the available optical mapping data from these two genomes to obtain deletions using BNG Solve and OptiDiff (with a segment length of 138kb). We then looked at the correspondence, defined as the percentage of overlap (in basepairs), between deletions found by BNG and OptiDiff and the sequence gaps. In Table 1 we show precision and recall values along with the *F*_1_-score (the harmonic mean of precision and recall), where the sequence gaps are used as the ground truth. Overlapping deletions are counted as true positives; false positives are deletions detected outside these overlapping regions and false negatives are sequence gaps which are not detected as deletions. The results demonstrate generally lower precision but far high recall for OptiDiff, resulting in higher *F*_1_-scores. This indicates a high correspondence between OptiDiff-detected deletions and gaps in the sequence alignments. The high level of agreement between these completely different approaches bolsters confidence in the OptiDiff results, and the high dissimilarity between the two accessions used demonstrate OptiDiff’s superior performance even across divergent genomes.

**Table 1:**
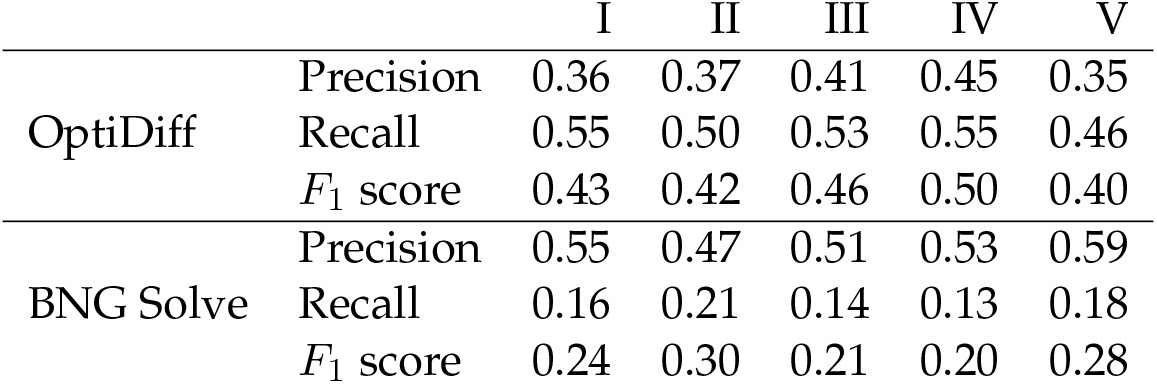
Correspondence between deletion sites detected by OptiDiff and BNG to gaps in genome alignments of Cvi to Col, in terms of *F*_1_-scores. The columns indicate the five chromosomes of *A. thaliana.*

|  |  | I | II | III | IV | V |
| --- | --- | --- | --- | --- | --- | --- |
| OptiDiff | Precision | 0.36 | 0.37 | 0.41 | 0.45 | 0.35 |
|  | Recall | 0.55 | 0.50 | 0.53 | 0.55 | 0.46 |
| | $F_1$ score | 0.43 | 0.42 | 0.46 | 0.50 | 0.40 |
| BNG Solve | Precision | 0.55 | 0.47 | 0.51 | 0.53 | 0.59 |
|  | Recall | 0.16 | 0.21 | 0.14 | 0.13 | 0.18 |
| | $F_1$ score | 0.24 | 0.30 | 0.21 | 0.20 | 0.28 |

### 3.6 OptiDiff is fast and accessible

Figure 11 depicts the time taken for SV detection with OptiDiff across different coverages for a 50Mb genomic region on a Linux server (Ryzen 7 3700X CPU) using 16 threads. Since OptiDiff does not include an assembly step, it is able to complete structural variation detection in below an hour for a chromosome of 50Mb with 120x coverage. In contrast, BNG Solve took over 10 times as long. OptiDiff is available as a command-line tool at https://github.com/akdel/OptiDiff. The code is written in Python with the help of the Numba library to increase performance in critical stages.

**Figure 11:**
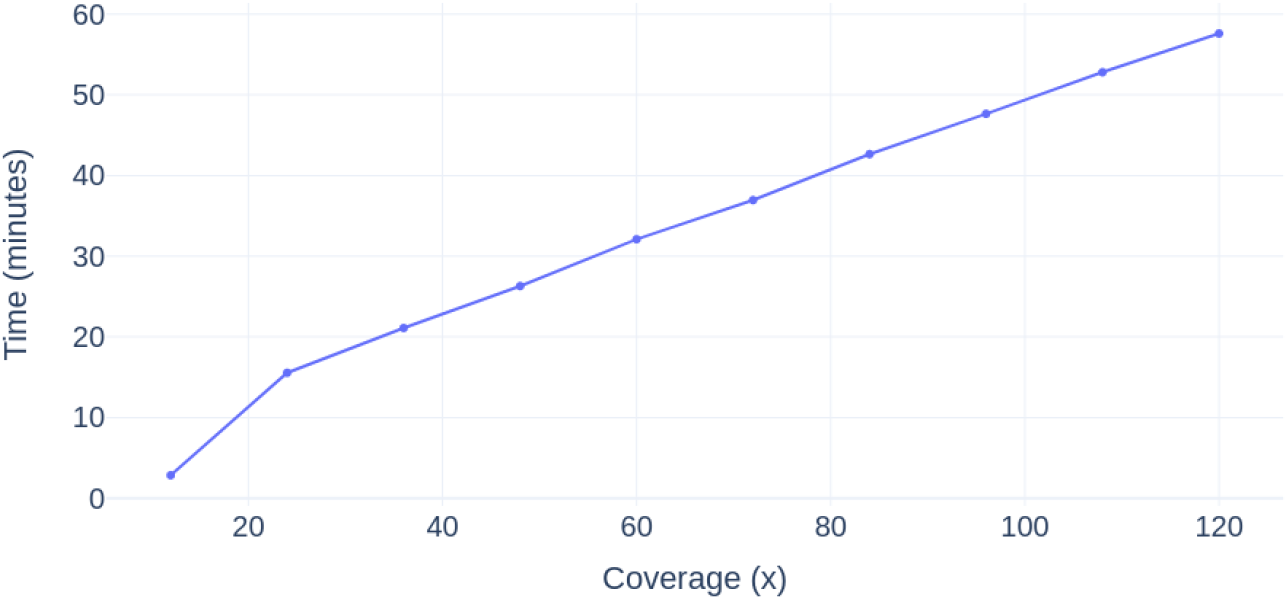
Time taken by OptiDiff to detect structural variations in a 50Mb genomic region across different coverages, using 16 threads.

## 4 Conclusion

We present OptiDiff, a tool for detecting and classifying structural variation sites from BNG optical mapping data. OptiDiff shows better detection performance than the state-of-the-art BNG Solve method in terms of precision, while maintaining comparable classification accuracies even at low coverages. OptiDiff shows high specificity and sensitivity across different SV types both in homozygous and heterozygous SV settings, and across highly similar and highly dissimilar genomes. On the *Arabidopsis* data, OptiDiff also shows greater overlap with other sequence-based technologies than BNG Solve.

Our method takes a single molecule approach, where each molecule is individually segmented and matched to an available reference assembly or genome map. The segment-matching gives the required information to detect the location of any type of SV using the change in coverage depths. Following this detection, OptiDiff uses the matching patterns of segments as evidence to classify SVs into deletions, duplications, translocations and inversions. A limitation of our approach is the inability to definitively classify SV types shorter than the predetermined segment length; these are labeled as short unspecific SVs. The approach used also makes it impossible to classify detected SV sites as insertions, since the inserted sequence is absent from the reference genome. We showed that different types of short SVs could be classified better with different segment lengths, indicating potential for an improvement to the algorithm where one could choose the optimal segment length for each SV type independently. An additional partial assembly step, where the molecules partially mapped to detected SV regions are extended to obtain the inserted maps, could help classify insertions. We leave these improvements for further investigation.

The future holds more advances in comparative genomics and in finding novel structural variants linked to phenotypes of interest. We expect OptiDiff and its extensions to play a role in this, as the advantages of using optical mapping in structural variation detection are becoming more evident.

## Acknowledgements

We thank Florian Jupe and Joseph Ecker for providing us with optical mapping data for *Arabidopsis thaliana* accessions; Aude Darracq, Geo Velikkakam James, Nikkie van Bers and Glenda Willems for fruitful discussions.

